# Microbial predator-prey interactions could favor coincidental selection of diverse virulence factors in marine coastal waters

**DOI:** 10.1101/439018

**Authors:** Etienne Robino, Aurore C. Poirier, Carmen Lopez-Joven, Jean-Christophe Auguet, Tristan P. Rubio, Chantal Cazevieille, Jean-Luc Rolland, Yann Héchard, Delphine Destoumieux-Garzon, Guillaume M. Charrière

## Abstract

Vibrios are ubiquitous in marine environments and opportunistically colonize a broad range of hosts. Strains of *Vibrio tasmaniensis* present in oyster farms can thrive in oysters during juvenile mortality events. Among them, *V. tasmaniensis* LGP32 behaves as a facultative intracellular pathogen of oyster hemocytes, a property rather unusual in vibrios. Herein, we asked whether LGP32 resistance to phagocytosis could result from coincidental selection of virulence factors during interactions with heterotrophic protists, such as amoeba, in the environment. To answer that question, we developed an integrative study, from the first description of amoeba diversity in oyster-farming areas to the characterization of LGP32 interactions with amoebae of the *Vannella* genus that were found abundant in the oyster environment. LGP32 was shown to be resistant to grazing by amoebae and this phenotype was dependent on previously identified virulence factors: the secreted metalloprotease Vsm and the copper efflux p-ATPase CopA. Using dedicated *in vitro* assays, our results showed that these virulence factors act at different steps during amoeba-vibrio interactions than they do in oysters-vibrio interactions. Hence, the virulence factors of LGP32 are key determinants of biotic interactions with multiple hosts ranging from protozoans to metazoans, suggesting that the selective pressure exerted by amoebae in marine coastal environments favor coincidental selection of virulence factors.

## INTRODUCTION

Vibrios are γ-proteobacteria living in aquatic environments ranging from saline to freshwater. They are ubiquitous in marine coastal environments and have evolved the capacity to colonize a broad range of hosts from protozoans to metazoans. Vibrios belong to the microbiota of healthy oysters but some species that behave as opportunistic pathogens can thrive in host tissues and cause disease ^1,2^. This occurs mostly as a result of environmental changes such as shifts in water temperature, exposure to high animal densities and stressful farming practices or upon immune suppression of host defenses by other microorganisms such as the OsHV-1 virus ^3,4^. Currently, vibrioses are recognized as a major factor limiting the development of aquaculture ^5^. In addition, vibrios can cause severe disease outbreaks in human populations, the best-known example being cholera. As they are multi-host pathogens, the ecology of vibrios depends on a serie of biotic interactions in the environment that influence their ecology, their transmission to animal and human hosts ^6,7^.

Such biotic interactions are also likely to favor the selection of virulence traits in vibrios. Predator-prey interactions are thought to generate some of the strongest forces driving a rapid co-evolution of both partners often referred as an arm-race ^8^. In natural environments, one of the main classes of bacterial predators are bacteriovorous protists. They can be found in most ecological niches inhabited by bacterial communities, and their predation activity through grazing share cellular and molecular processes with metazoan immune cells antibacterial responses, in particular phagocytosis ^9^. Hence, bacterial predation by protists is thought to favor co-incidental selection of bacterial virulence factors in opportunistic pathogens having multiple hosts (generalists) ^10^.

Although vibrios have long been considered as extracellular pathogens, an increasing number of studies have demonstrated that some vibrios can adopt intracellular stages ^11–15^. Strains of *Vibrio tasmaniensis,* which are present in the water column and sediment of oyster farms and can thrive in oyster flesh during juvenile mortality events ^4,16^, have adopted intracellular stages needed for pathogenicity ^17^. Thus, the oyster pathogen *V. tasmaniensis* LGP32 isolated from moribund oysters ^18^, enters and survives in oyster immune cells, the hemocytes ^17^; this intracellular stage being required for its virulence in oysters and leading to cytotoxic damages ^17,19^. Some virulence factors of *V. tasmaniensis* LGP32 have been described. They include secreted proteases as Vsm ^20^ or Vsp ^21^, antioxydants such as the superoxide dismutatase SodA, and efflux pumps to detoxify heavy metals such as the copper p-ATPase CopA ^19^.

Herein, we asked whether predator-prey interactions in microbial communities could represent a driving force that selects for virulence and resistance traits favoring the emergence of facultative intracellular pathogens among vibrios. For that, we studied the interactions between LGP32 and environmental free-living marine amoebae. After describing amoeba diversity in oyster-farming areas, we characterized LGP32 interactions with amoebae of the *Vannella* genus, which belongs to the most ubiquitous taxonomic group in our environmental survey. Our results revealed that LGP32 resists to the grazing by the environmental isolate *Vannella sp*. AP1411. Through a combination of flow cytometry, confocal and electron microscopy analyses, we showed that LGP32 delays intracellular traffic and food vacuoles formation in amoeba. By analyzing multiple mutant strains deleted for previously characterized virulence factors in oysters, we found that the copper p-ATPase efflux pump CopA and the secreted metalloprotease Vsm are both required for the resistance of LGP32 to grazing by marine amoebae. CopA appears to play a role in the intracellular survival of LGP32 whereas Vsm appears to play a role in the inhibition of amoebae migration on during grazing. Altogether, our results indicate that virulence factors of opportunistic pathogens can be involved in different types of interactions with multiple hosts from protozoans to metazoans, and suggests that the selective pressure exerted by heterotrophic protists in the oyster environment can favor co-incidental selection of virulence factors and pathogen emergence.

## Experimental procedures

### Bacterial strains and growth conditions

*E.coli* strain SBS363 was grown in Luria-Bertani (LB) or LB-agar (LBA) at 37°C. The marine amoeba *Vannella* sp. AP1411 (isolated in this study) was grown in 70% sterile seawater (SSW) with *E. coli* wild-type strain SBS363 at 18°C, for 3 days prior experiments. Vibrios strains used in this study are *V. tasmaniensis* LMG20012^T^, *V. tasmaniensis* LGP32 and previously described LGP32 deletion mutants **Δ***copA*, **Δ***cusAB*; **Δ***sodA*; **Δ***vsm*, **Δ***vsp*, **Δ***inhA* ^19–21^. Vibrios strains carrying the pMRB-GFP plasmid were grown in LB + NaCl 0.5M supplemented with chloramphenicol (10μg mL^-1^) at 20°C, for 24 hours prior experiments.

### Isolation of environmental amoebae from the Thau Lagoon

Water, oysters and sediment from the Thau lagoon (South France) were sampled at least two times per season between 2014 and 2015. Water was collected next to oyster tables at the Bouzigues station Ifremer-REPHY (GPS: N 43°26.058′ E 03°39′.878’) and filtered, on the boat, with a 180 μm pore size nylon filter. In the lab, the water was re-filtered using a 8.0 μm pore size MF-Millipore membrane. The 8.0 μm pore size membrane was then cut in four pieces and each quarter was put upside down on a lawn of *E. coli* SBS363 seeded on SSW-agar. Oyster gills were cut in 4 pieces of roughly 0.5 cm^2^ and each quarter was put upside down on a lawn of *E. coli* SBS363 seeded on non-nutrient SSW-agar. Sediment was collected under the oyster tables (depth of 9 meters) by core sampling. One gram of sediment was then deposited in the center of a lawn of *E. coli* SBS363 seeded on SSW-agar, in triplicate. After 1 to 3 weeks, depending of the sampling season, migrating amoebae were observed at the periphery of the plates. An agar square of 0.5×0.5 cm containing a single amoeba was then cut and deposited upside down on a fresh lawn of *E. coli* SBS363 seeded on SSW-agar. This was repeated several times to isolate and replicate amoebae. DNA was extracted using the High Pure PCR Template Preparation Kit (Roche), according to the manufacturers’ protocol.

### Identification of isolated marine amoebae from Thau Lagoon

In order to infer the phylogenetic assignation of environmental amoebae, the v7 region of the 18s rDNA gene was sequenced using specific primers. The primers were designed using SILVA database dedicated tool (http://www.arb-silva.de/search/testprime). The selected primers, Amo_AP_1154_F: 5’ GAGRAAATTAGAGTGTTYAAAG 3’ and Amo_AP_1470_R: 5’TTATRGTTAAGACTA CGACGG 3’, were used to amplified the hypervariable region V7 of 18S rDNA gene at an annealing temperature of 54°C. PCR amplicons were cloned using the TOPO TA Cloning Kit (Invitrogen), according to the manufacturers’ protocol. Nucleotides sequences were determined by Sanger sequencing (GenSeq platform, Labex CEMEB, Montpellier, France). Sequence homologies were searched using BLASTn available at the NCBI web site (National Centre for Biotechnology Information ^22^ (Table S1). For phylogenetic analyses, sequences were aligned in MOTHUR ^23^ and imported into ARB software ^24^ loaded with the silva database (http://www.arb-silva.de). A base frequency filter was applied to exclude highly variable positions before adding sequences to the maximum parsimony backbone tree using the parsimony quick add marked tool implemented in ARB, thereby maintaining the overall tree topology provided by default. In order to verify that our Sanger sequencing strategy allowed us to identify the entire diversity of amoebae that could be isolated through culturing technics. The total DNA from our different amoebae cultures was analyzed by performing barcoding on the v7 region of the 18S rDNA gene using generic primers that allowed amplifying more than 80% of all the eukaryote 18s rDNA sequences available in SILVA database (F-v7-1173: CCT GCG GCT TAA TTT GAC and R-v7-1438: CAT CAC AGA CCT GTT ATT GC). Sequencing was performed by Genome Quebec facility, Montreal, Canada.

### Grazing assay

To prepare the co-culture of vibrios and amoebae, 1 mL of vibrio overnight culture (3.10^9^ bacteria. mL^-1^) was mixed with 100 μL of three days old *Vannella* sp. AP1411 culture (5.10^5^ cells. mL^-1^) or with 100 μL of 70% SSW for control condition. A volume of 50μL per well of the mixed culture were seeded on top of 500 μL of 1% SSW-agar, in 24-wells plate with transparent flat bottom. Amoebae and bacteria lawn were carefully homogenized in the wells and let dry, during 4 hours at room temperature under the flow a sterile hood, and then incubated at 18 °C in a humidified atmosphere. GFP fluorescence intensity was measured every day over 7 days using a TECAN plate reader (λex 480 nm/λem 520 nm). Then, to estimate the effect of the amoebae grazing activity on the abundance of living vibrios expressing GFP, the fluorescence intensity of the wells containing amoebae was compared to the fluorescence of vibrios lawn without amoebae, and express as a ratio. Each condition was performed in technical triplicates and the results shown are the average of three independent experiments. Error bars represent the standard error of the mean (±SEM). Statistical analysis was performed using RM-ANOVA over the independent experiments.

### Monitoring amoeba growth and association with GFP-vibrios

To estimate the proliferation of *Vannella sp*. AP1411 at days 1, 3 and 6 in grazing assays, amoebae were directly pictured on top of the agar under phase light and manually counted in triplicate for each condition. Then, cells were flushed from the SSW-agar surface with 1 mL of 70% SSW and fixed during 30 minutes with 2% paraformaldehyde at room temperature. Pellets were washed and suspended in 500μL of PBS for flow cytometry analysis (BD FACS Canto). Data analyses were performed using Flowing Software by gating on amoeba cells using SSC/FSC parameters then quantifying the percentage of GFP+ amoebae (as shown in Figure S2). In order to estimate the phagocytosis index, which corresponds to the average number of phagocytosed bacteria per cell, the mean intensity of GFP fluorescence per cell was normalized on the mean fluorescence per GFP+ bacteria (as shown in Figure S3). The experiments were performed with three technical replicates per condition in each experiment, and depicted results are representative of at least two independent experiments. Error bars represent the standard error of the mean (± SEM). Statistical analysis was performed using two ways ANOVA with Holm-Sidak’s multiple comparisons test.

### Microscopy

The different amoeba shapes were observed on living amoebae using phase contrast (Inverted wide-field epifluorescence microscope, Zeiss). A culture of amoebae in 4 wells plates was washed twice with 70% SSW before observation. The shape of the floating forms was recorded by scraping in the cell suspension followed by immediate observation. The cystic form was observed after amoeba starvation for a week. Intracellular organization of the amoeba was observed using transmission electron microscopy. Briefly, amoebae cultured in flask were washed twice using 70% SSW, and fixed in 2.5% glutaraldehyde for 2 hours in the dark at room temperature, then overnight at 4°C. After centrifugation, the supernatant was replaced by 3% low melting-point agarose (Sigma) and the pellet was resuspended. Cells entrapped in 3% agar were immersed in a solution of 2.5% glutaraldehyde in PHEM buffer (1X, pH 7.4) overnight at 4°C. Then they were rinsed in PHEM buffer and post-fixed in a 0.5% osmic acid for 2 h in the dark and at room temperature. After two rinses, samples were dehydrated in a graded series of ethanol solutions (30-100%). Samples were embedded in EmBed 812 using an Automated Microwave Tissue Processor for Electronic Microscopy, Leica EM AMW. Ultrathin sections (70nm; Leica-Reichert Ultracut E) were collected at different levels of each block, then counterstained with uranyl acetate 1.5% in 70% Ethanol and lead citrate and observed using a Tecnai F20 transmission electron microscope at 120KV in the CoMET MRI facilities, INM, Montpellier France.

Confocal fluorescence microscopy was performed on amoebae suspensions after PFA fixation to observe the localization of intra-amoeba vibrios. Cell suspensions were cytospun on glass slides for 10 min at 800 g. Glass slides were then stained with 0.25 μg mL^-1^ DAPI (Sigma) and 0.1 μg mL^-1^ Blue Evans (Sigma). Confocal fluorescence imaging was performed using a 63X oil objective and images were captured using a Leica TCS SPE confocal scanning laser microscope, at 1 airy to ensure the focal plan.

### Time-lapse imaging and cell tracking

Co-cultures of vibrios and amoebae were prepared following the same protocol than for grazing assays. Time-lapse imaging started 24 hours after the beginning of the grazing assay. Acquisitions were obtained using Inverted wide-field epifluorescence microscope (Zeiss) and B&W coolsnap camera. Time-lapse were done during 30 minutes with 1 frame every 30 seconds. TrackMate Tools software (ImageJ) was used for manual tracking to determine velocity and migration distance by amoebae on different vibrios lawns (Supplementary Movie 1, 2 and 3). Migration distances were measured for 18 amoebae at the same time (i.e. 20 frames by amoebae trace) for each condition. Each track depicted corresponds to the migration of an amoeba during 10 minutes. The migration path was determined by the x and y positions of amoebae tracked at each frame. Depicted results are representative of at least two independent experiments. Velocities were measured for 16 amoebae tracks, only tracks made of at least 5 frames were considered. Each result presented corresponds to the average velocity of one amoeba per track. Depicted results are representative of at least two independent experiments. Error bars represent the standard deviation (± SD). Statistical analysis was performed using Kruskal-Wallis test with Dunn’s multiple comparisons test.

## Results

### Marine amoeba of the *Vannellidae* order are the main cultivable free-living amoebae isolated from the oyster-farming environment

Although amoebae represent an important category of benthic grazers of bacterial communities, their diversity remains poorly explored in marine environments, particularly in Mediterranean lagoons, which are exploited for oyster farming. Therefore first objective of this study was to identify and isolate the most prevalent amoebae in marine environments hosting oyster cultures, *i.e.* grazers that can exert permanent grazing pressure on vibrios pathogenic for oysters. Amoebae diversity was here characterized in the Thau lagoon, in the vicinity of oyster farms. Amoebae were isolated from the water column, oyster gills and sediment at all seasons between the year 2014 and 2015, and at least two cultures of amoebae per fractions and per seasons were analyzed. Cloning-sequencing of the v7 hypervariable region of 18S rDNA gene was used to identify the isolated amoebae. A total of 110 sequences were analyzed. From 24 different cultures, 103 amoebae sequences showed more than 90% of sequence homologies with the *Vannella* genus, which belongs to the *Vannellidae* order. The 103 sequences were classified into 11 different Operational Taxonomic Units (OTUs) (Figure 1 and Table S1). In parallel, barcoding of the different amoebae cultures confirmed that only species belonging to the *Vannella* genus were present in the samples (Supplemental Figure S1). One OTU of *Vannella sp.* referred to as AP1411 was ubiquitous, being found in the 3 fractions and at the four seasons (arrow in Fig. 1). It was therefore chosen to study further amoebae-*Vibrio* interactions. The trophozoite, the planktonic and the cystic forms were typical of the *Vannella* genera and very similar to *Vannella plurinucleolus* ^25^, which was also the closest 18S matching sequence for this isolate (98% Blastn score) (Figure 2A). Indeed, transmission electron microscopy of trophozoïtes revealed a typical intracellular organization of *Vannella* with a well-defined, thick plasma membrane, one nucleus (sometime two or three), numerous mitochondria and a rich vacuolar system with large digestive vacuoles containing remnants of degraded bacteria, and some phagosomes containing individual bacteria (Figure 2B).

**Fig. 1.**
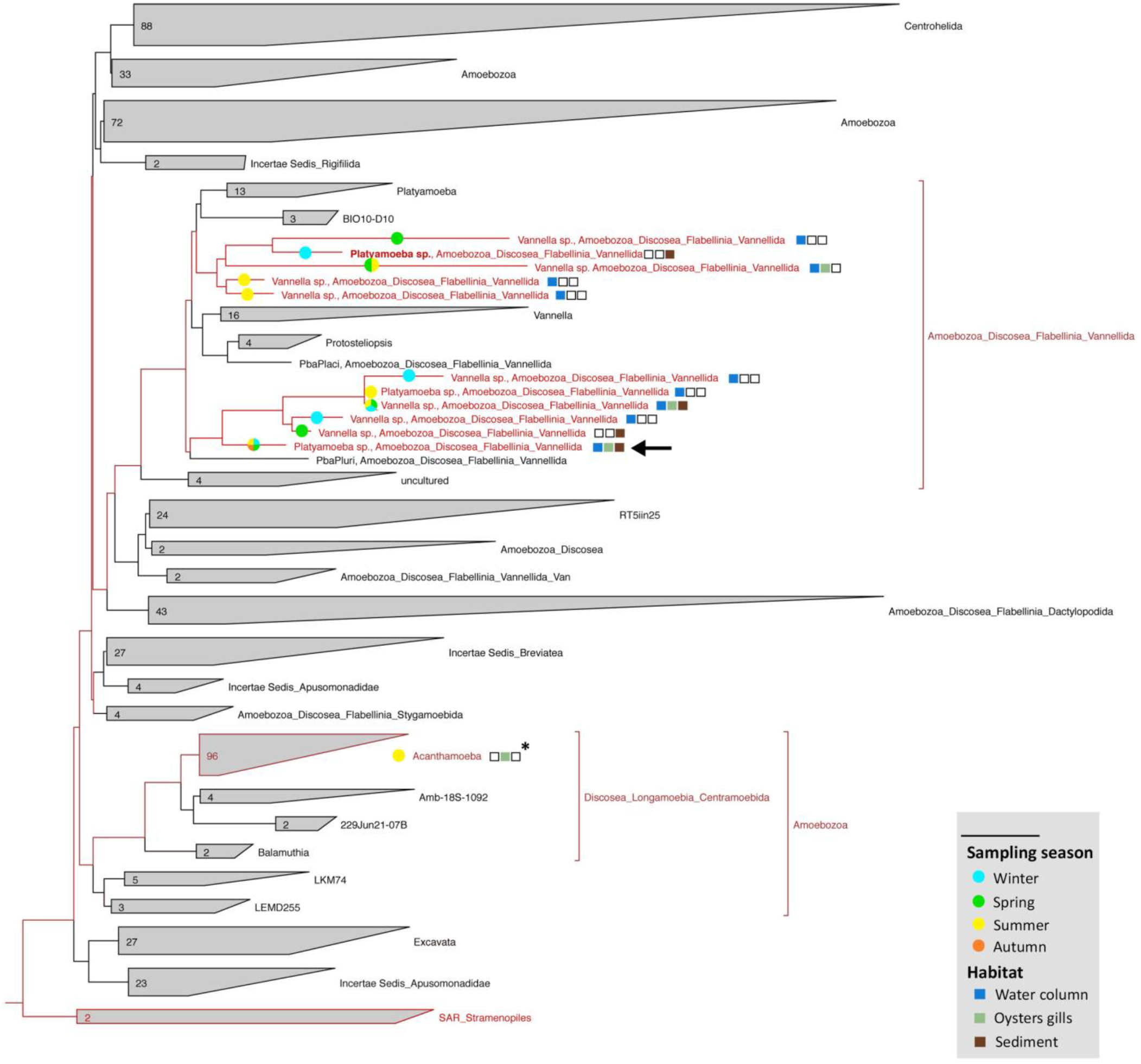
Most of the isolated free-living amoebae from the Thau lagoon belong to Vannellidae order. Phylogenetic 18S rRNA-based trees of the Amoebozoa constructed with ARB software (Ludwig et al., 2004; http://www.arb-home.de) loaded with the silva database (http://www.arb-silva.de). A base frequency filter was applied using the parsimony quick add marked tool implemented in ARB. Our isolates are written in red. Most of the isolated amoebae during the environmental survey from water column (blue squares) and/or the sediment (brown squares), and/or oyster gills (light green squares) during winter (light blue circles) and/or spring (green circles) and/or summer (yellow circles) and/or autumn (orange circles) belong to the *Vannellidae* order.

**Fig. 2.**
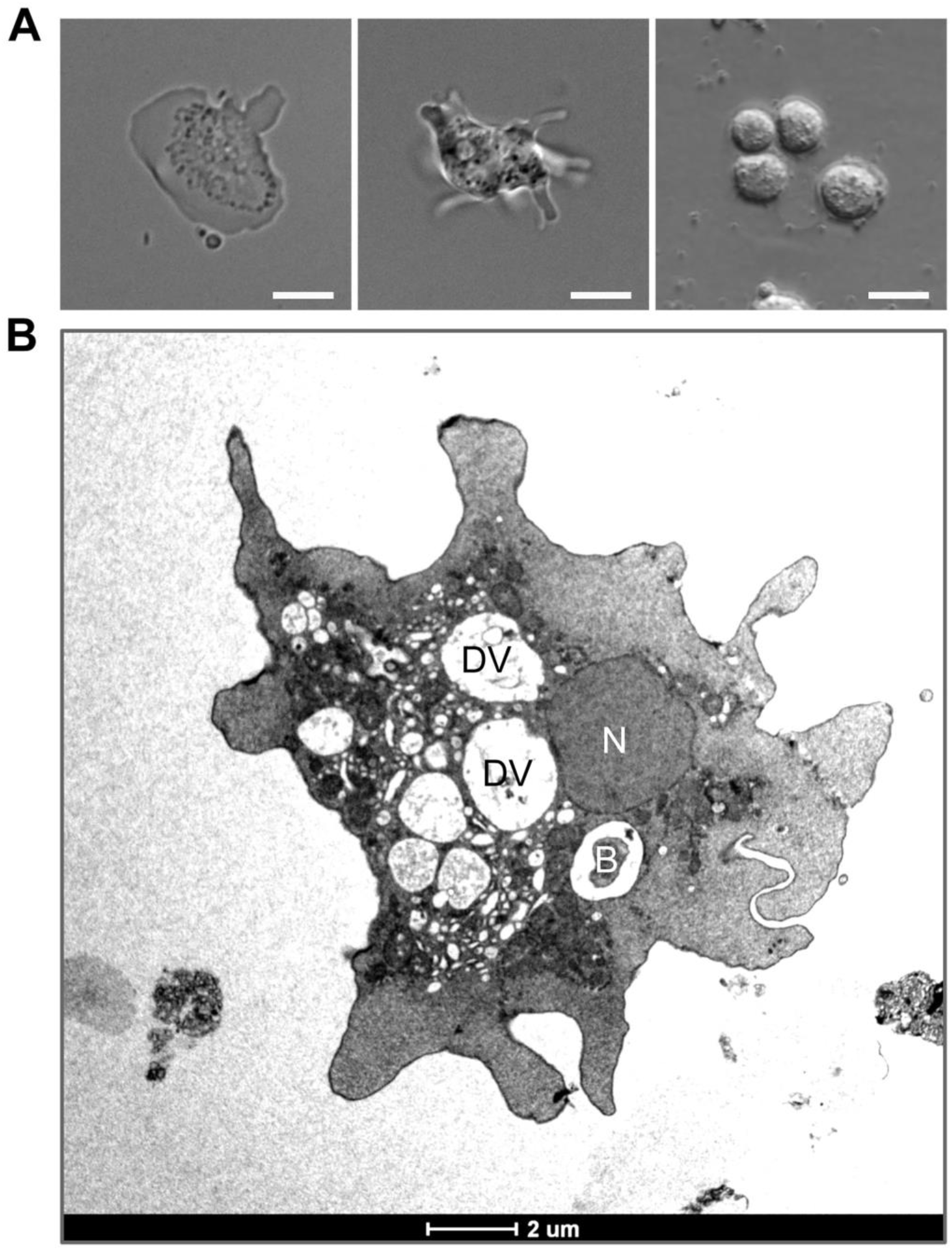
*Vannella sp*. AP1411 isolated from Thau lagoon, France. **(A)** Different forms of *Vannella* sp. AP1411 under phase light microscopy. Trophozoite (left), pelagic (center) and cystic form (right). Scale bar : 10 μm. **(B)** Transmission electron microscopy of trophozoite shape of the amoeba *Vannella* sp. AP1411. Whole cell overview showing nuclei (N) and several digestive vacuoles (DV) with or without *E. coli* SBS363 bacteria inside (B). Scale bar: 2 μm.

### The pathogen LGP32 resists to grazing by *Vannella* sp. AP1411

To study the interactions between vibrios and *Vannella* sp. AP1411, grazing assays were performed on solid media (non-nutrient seawater agar). *Vibrio* strains (pathogenic or non-pathogenic to oysters) that constitutively express the Green Fluorescent Protein (GFP) were used to seed the plates and monitor the abundance of live vibrio cells by fluorescence quantification. *Vannella sp*. AP1411 growth was also monitored in parallel, by counting the cell density at the surface of the grazing lawns. When amoebae were cultured in the presence of the non-virulent *Vibrio tasmaniensis* LMG20012^T^, a rapid decay of GFP fluorescence was observed 4 days after the beginning of the experiment; whereas in the case of the pathogenic *V. tasmaniensis* LGP32, the amount of GFP-expressing bacteria remained significantly more stable for a longer period of time (Figure 3A). Amoebae growth mirrored the kinetics of the GFP fluorescence decay of the vibrios. Indeed amoebae density increased significantly faster in the presence of LMG20012^T^ than in the presence of LGP32 (Figure 3B). These data indicate that LGP32 does not support well *Vannella sp*. AP1411 growth, whereas LMG20012^T^ is more favorable.

**Fig. 3.**
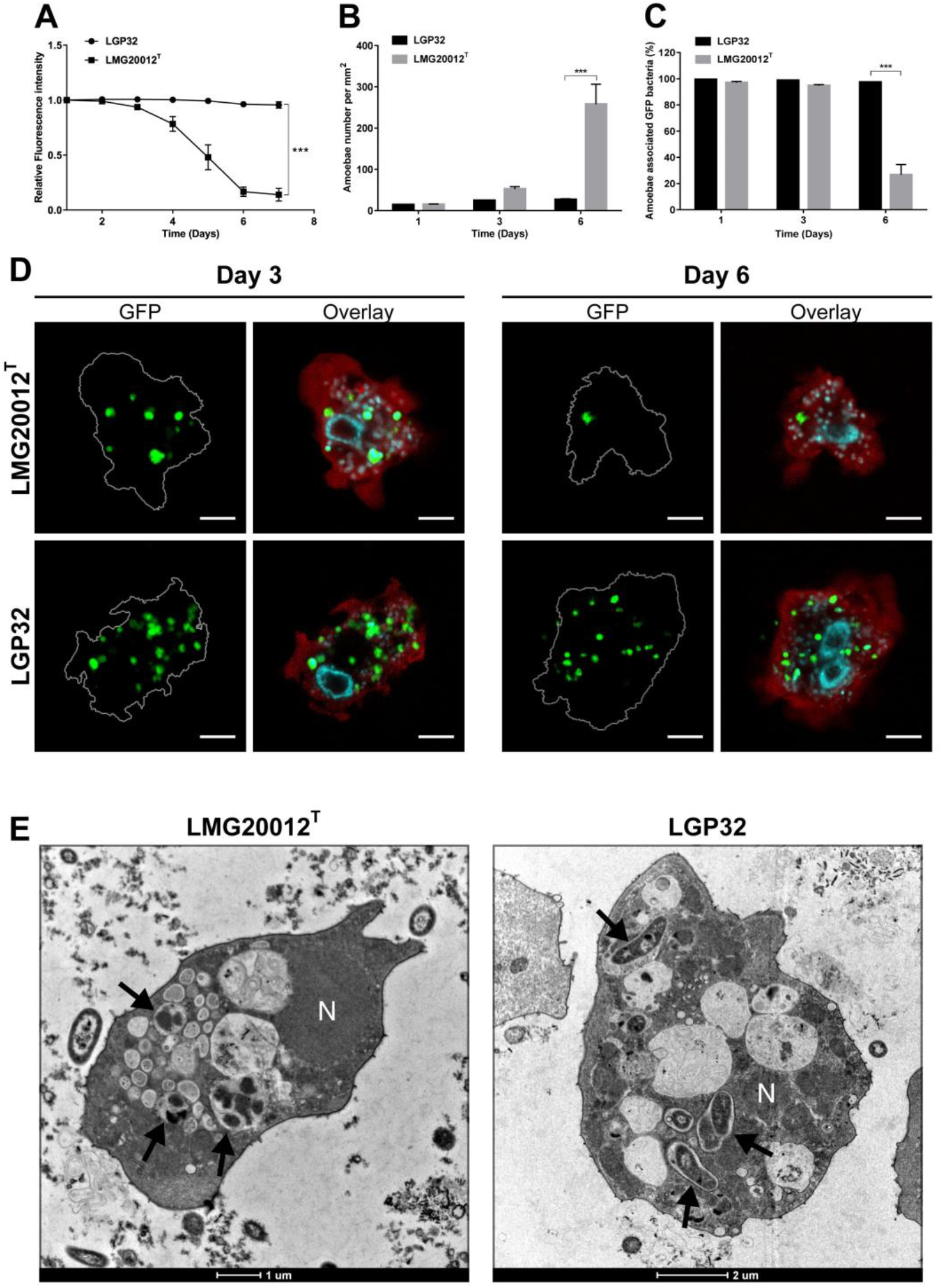
LGP32 is resistant to grazing by seawater amoebae *Vannella sp*. AP1411. Amoebae were infected with GFP mutants of LGP32 or of the avirulent LMG20012^T^. **(A)** LGP32 is more resistant to grazing than the avirulent control LMG20012^T^. Bacterial resistance was assessed by monitoring the intensity of the bacterial GFP fluorescence. Each condition was performed in triplicate and the results represent the average of three independent experiments. Values are presented ± SEM *** P < 0.001 (RM-ANOVA). **(B)** LGP32 disadvantages amoebae growth compared to the avirulent control LMG20012^T^. Amoebae growth was monitored by manual counting under phase light microscopy. Each condition was performed in3 technical replicates and depicted results are representative of 2 independent experiments ± SEM. Data were analyzed by two-way ANOVA with Holm-Sidak’s multiple comparisons test, *** P < 0.001. **(C)** LGP32 persists within amoebae for 6 days. The number of amoebae associated with GFP bacteria fluorescence was measured by flow cytometry. Each condition was performed in 3 technical replicates and depicted results are representative of 2 independent experiments ± SEM. Data were analyzed by two-way ANOVA with Holm-Sidak’s multiple comparisons test, *** P < 0.001. **(D)** LMG20012^T^ tends to cluster within amoebae but not LGP32. Amoebae withdrawn from co-cultures of amoebae and bacteriae were observed by confocal microscopy 3 days and 6 days after contact. LMG20012^T^ was rapidly degraded and forms aggregates from day 3 after infection. LGP32 is less degraded than LMG20012^T^ and forms spots at day 3 and 6. Nuclei were stained with DAPI and amoebal proteins with Blue Evans. Scale bar: 5 μm. **(E)** Transmission electron microscopy was performed on amoeba 6 days after grazing on LGP32 or LMG20012^T^. Avirulent LMG20012^T^ vibrios were found clustered in intracellular vacuoles with an altered shape and appearance (arrows), compared to extracellular vibrios around the amoeba. Virulent LGP32 vibrios were found mostly in individualized phagosomes, and appeared intact (arrows). Scale bar: 1 μm and 2 μm.

To determine whether the amoeba growth defect observed in the presence of LGP32 was due to a higher resistance to grazing of the pathogenic LGP32 strain, we next compared the capacity of *Vannella* sp. AP1411 to phagocytose LGP32 and LMG20012^T^. After three days of grazing, 100% of amoebae were similarly associated with GFP-LMG20012^T^ or GFP-LGP32, as revealed by flow cytometry. However at day 6, the percentage of amoebae associated with GFP-LMG20012^T^ decreased whereas it remained stable with GFP-LGP32 (Figure 3C). Most of the vibrios (both GFP-LGP32 and GFP-LMG20012^T^) were found inside amoebae as revealed by confocal microscopy (Figure 3D), showing that vibrios were readily internalized and not just adherent to the cell surface of amoebae. Interestingly, at day 3, GFP-LMG20012^T^ clustered inside cytoplasmic vacuoles (3 to 5 bacteria per confocal sections per vacuole), whereas very little intracellular clusters of GFP-LGP32 were observed. At day 6, most amoebae were devoid of GFP-LMG20012^T^ as revealed by flow cytometry quantification, suggesting that they had been killed and digested. In contrast, GFP-LGP32 were still intact inside amoebae and not much clustered.

To investigate further the intracellular fate of the vibrios inside amoebae, transmission electron microscopy was performed at day 3 (Figure 3E). Most LMG20012^T^ cells were clustered inside digestive vacuoles and showed an altered morphology, as expected during the process of food vacuole formation. In contrast, only few LGP32 cells were observed inside amoebae; when ingested, they remained in individual phagosomes; their morphology appeared preserved. Altogether these results show that LGP32 resists to the grazing activity of *Vannella sp*. AP1411 as opposed to the non-virulent LMG20012^T^ and suggest that resistance to intracellular degradation relies on the inhibition of the intracellular trafficking during phagosome maturation and food vacuole formation.

### The Vsm and CopA virulence factors participate in LGP32 resistance to grazing by marine amoebae

We tested here whether mechanisms of virulence and resistance to host defenses evidenced in the interaction with oysters could also play a role in LGP32 interaction with environmental amoebae. On the one hand, we examined stress resistance systems, i.e. the antioxidant SodA, and the copper resistance systems CopA and CusAB, involved in intraphagosomal resistance to antimicrobial activities of oyster immune cells ^19^. On the other hand, we examined secreted virulence factors, i.e. the metalloprotease Vsm and the serine protease Vsp, that both play a role as virulence factors in oyster pathogenesis ^20,21^, as well as the metalloprotease InhA (the ortholog of PrtV) that has been involved in multiple biotic interactions of vibrios ^26^. Among the six isogenic strains of LGP32 tested, only the **Δ***copA* and **Δ***vsm* deletion mutants were significantly more susceptible to grazing than the wild type strain, as shown by a faster clearance of GFP-vibrios on grazing lawns (Figure 4A). On lawns of both **Δ***copA* and **Δ***vsm*, rapid bacterial clearance correlated with fast amoeba growth; amoeba density was significantly higher than on wild-type LGP32 lawns (Figures 4B and 4D). These results show that both CopA and Vsm play a role in the resistance of LGP32 to grazing by *Vannella sp*. AP1411.

**Fig. 4.**
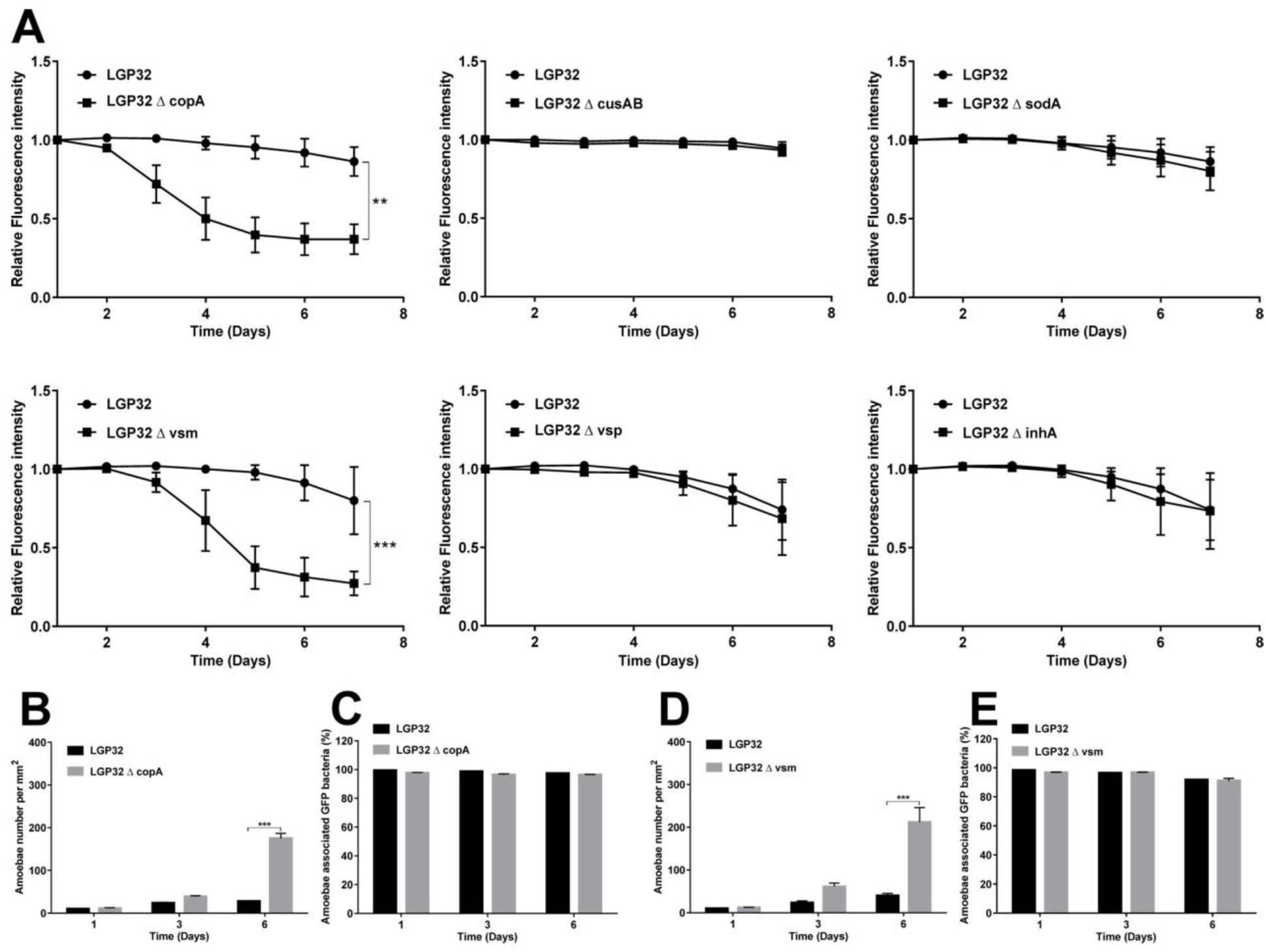
Copper p-ATPase efflux pump CopA and metalloprotease Vsm are involved in LGP32 resistance to grazing by *Vannella sp.* 1411. **(A)** *∆copA* and *∆vsm* mutant strains are more sensitive to grazing by amoeba but not *∆cusAB, ∆sodA, ∆vsp, and ∆inhA* strains. Bacterial resistance was assessed by measurement the fluorescence of the GFP-expressing bacteria, after contact with amoebae. Each condition was performed in triplicate and the results shown are the average of three independent experiments. Values are presented ± SEM, ** P < 0.01, *** P < 0.001 (RM-ANOVA). **(B-D)** *∆copA* and *∆vsm* mutant strains promote amoebal growth compared to the wild-type LGP32. Amoeba growth was monitored by manual counting under phase light microscopy. Each condition was counted in three technical replicates. The results shown are representative of two independent experiments ± SEM. Data were analysed by two-way ANOVA with Holm-Sidak’s multiple comparisons test, *** P < 0.001. **(C-E)** The percentage of phagocytosis is not affected in *∆copA* and *∆vsm* mutant strains compared to the wild-type LGP32. In two independent experiments, number of amoebae associated with GFP bacteria was measured by flow cytometry. Each condition was performed in three technical replicates. The results shown are representative of two independent experiments ± SEM. Data were analyzed by two-way ANOVA with Holm-Sidak’s multiple comparisons test.

### The p-ATPase CopA confers resistance to intracellular killing whereas the extracellular protease Vsm inhibits amoebae motility during grazing

To investigate further the role of LGP32 virulence factors in resistance to amoeba grazing, we analyzed different steps of the grazing process, from phagocytosis to intracellular degradation. Flow cytometry analyses showed that the percentage of phagocytosis (Figures 4C and 4E) and the phagocytosis index (Figure S3) were not significantly different from the wild-type LGP32, which suggested that the efficiency of the phagocytosis process was not dependent on CopA or Vsm. Confocal and TEM microscopy, showed that as for wild-type LGP32, internalized **Δ***vsm* mutants were mostly observed as isolated intact cells inside individual phagosomes within amoeba cytoplasmic space. In contrast, **Δ***copA* mutants behaved as the non-virulent LMG20012^T^, clustering inside digestive vacuoles and showing an altered morphology at day 6 (Figure 5A-B).

**Fig. 5.**
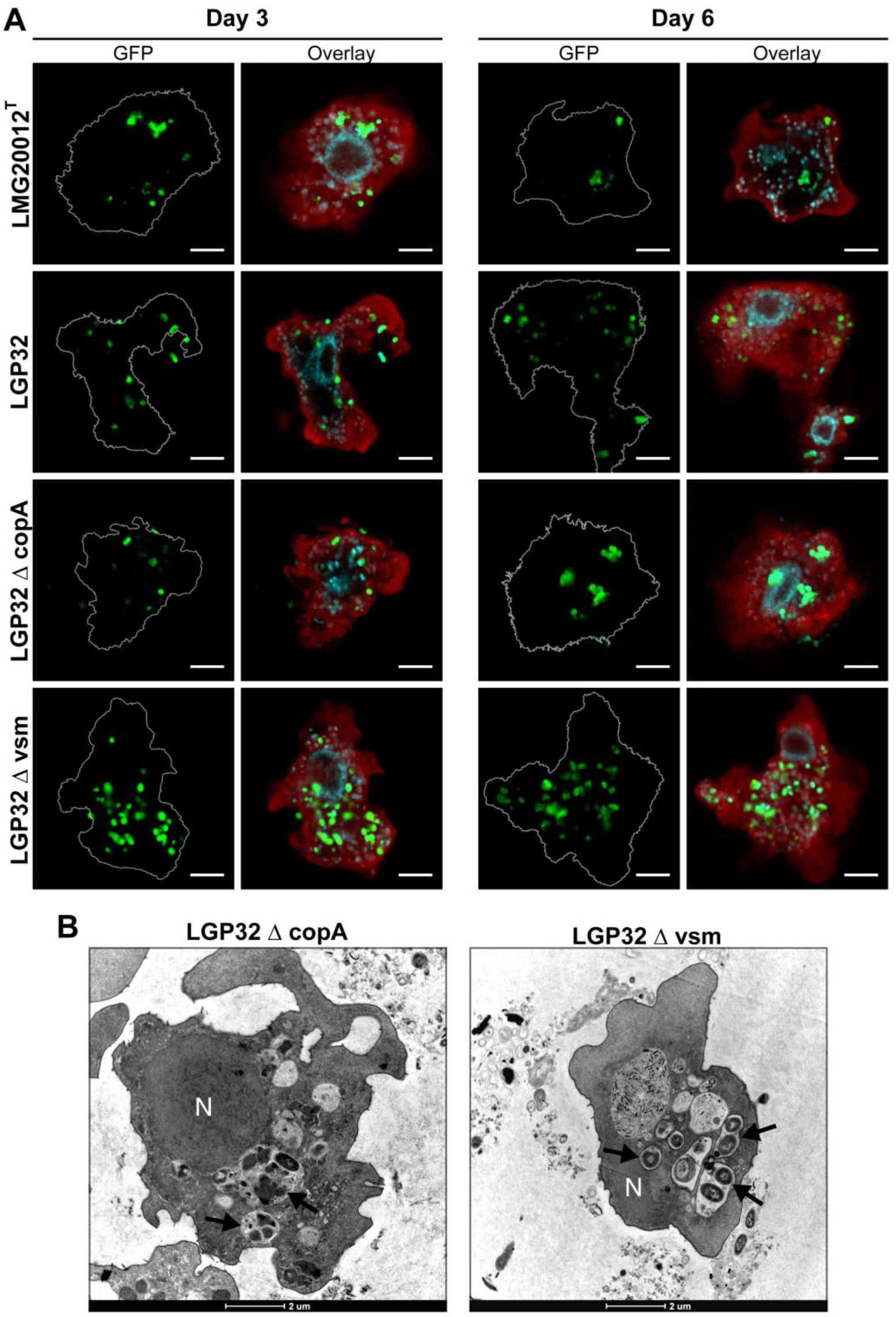
Copper p-ATPase efflux pump CopA is necessary to resist to intracellular degradation in amoeba whereas the metalloprotease Vsm is not. **(A)** Intracellular vibrios were observed 3 days and 6 days after infection, by confocal microscopy. All strains were phagocytosed but only the *∆copA* mutant strain forms aggregates within amoebae in a similar manner than the avirulent strains LMG20012^T^. Nuclei were stained with DAPI and amoeba proteins with Blue Evans. Scale bar: 5 μm. **(B)** Transmission electron microscopy was performed on amoeba 6 days after grazing on *∆copA* or *∆vsm* mutant strains. Mutant *∆copA* vibrios were found clustered in intracellular vacuoles with an altered shape and appearance (arrows), like the avirulent LMG20012^T^ strain. Mutant *∆vsm* vibrios were found mostly in individualized phagosomes, and appeared intact (arrows), like the LGP32 wild-type strain. Scale bar: 2 μm.

As the intracellular fate of Δ*vsm* did not differ from the wildtype LGP32, the role of Vsm in the resistance of LGP32 to grazing was investigated further. Since Vsm is a metalloprotease secreted extarcellularly, we asked whether it could alter the grazing behavior of *Vannella sp*. AP1411. The trajectory of amoebae was similar on lawns of wild-type LGP32, **Δ***vsm* and **Δ***copA*, as determined by time-lapse microscopy (Figure 6A, B and C). However, migratory speed was remarkably accelerated by ~ two fold (p<0.001) on the **Δ***vsm* grazing lawn compared to the wild-type and the **Δ***copA* grazing lawns (Supplementary Movies 1, 2 and 3) with mean velocities of 8.5, 3.9 and 3.8 microns per minute for **Δ***vsm,* wild-type LGP32 and **Δ***copA,* respectively (Figure 6D). These results indicate that the metalloprotease Vsm acts extracellularly by inhibiting amoeba motility, which likely disturbs amoeba grazing efficiency, whereas CopA confers intravacuolar survival capacity required for altering the normal formation of the digestive vacuole.

**Fig. 6.**
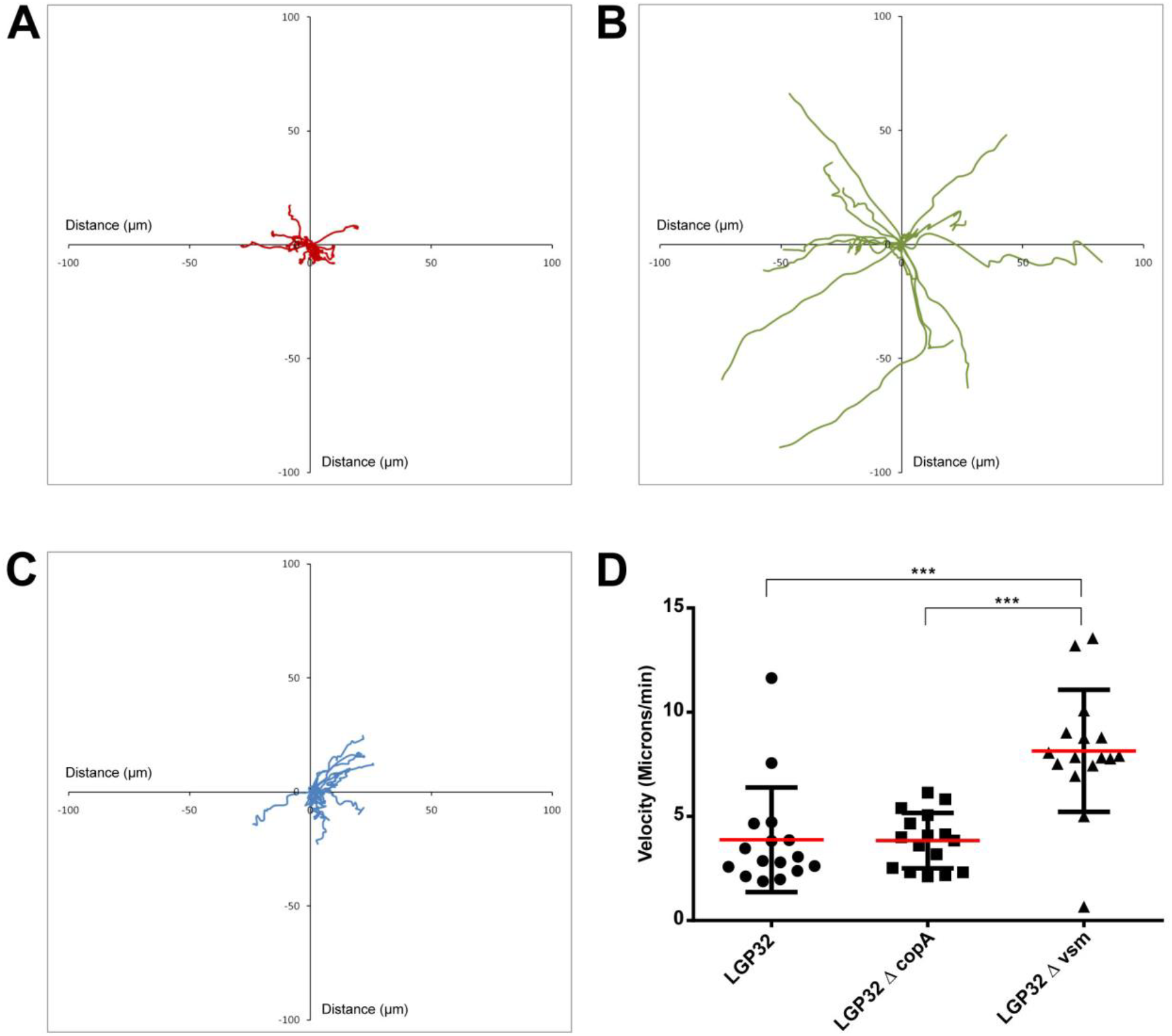
Extracellular metalloprotease Vsm decreases the velocity of the amoebae *Vannella sp*. AP1411 and alters amoeba grazing efficiency. Amoebae migration on LGP32wt **(A)**, ∆*vsm* **(B)** or ∆*copA* **(C)** mutant strains lawns during 10 minutes. Each presented track corresponds to the migration of one amoeba. The results shown are representative of at least two independent experiments. **(D)** Average velocity of amoebae on LGP32wt, ∆*copA* or ∆*vsm* mutant strains. The results are representative of at least two independent experiments ± SD. Data were analyzed by Kruskal-Wallis test with Dunn’s multiple comparisons test, *** P < 0.001.

## Discussion

In the present study, we investigated whether interactions between environmental phagocytes, i.e. amoebae, and vibrios could participate to the co-incidental selection of virulence factors in marine coastal waters. For that, we performed comparative interaction studies between the oyster pathogen *Vibrio tasmaniensis* LGP32 or LGP32 isogenic mutants attenuated for virulence in oysters and amoebae isolated from the wild. By using an amoeba isolate of the *Vannellidae* order, which we found to be ubiquitous in oyster farms of the Thau lagoon, we showed that *V. tasmaniensis* LGP32 is resistant to the grazing activity of *Vannella sp*. AP1411; in a similar manner than it resists to killing by oyster phagocytes, by involving the virulence factors CopA and Vsm.

Amoebae of the *Vannellidae* order were found here to be ubiquitous in the Thau lagoon, a Mediterranean coastal lagoon used for oyster farming (20% of its total surface). Indeed, most of the isolated amoebae branched into the *Vannella* genus, independently of seasons. Together with the present study, the current literature suggests Vannellidae forms a rather abundant order of amoeba in aqueous environments, particularly in saltwater, but they have also been frequently isolated from freshwater ^25^. As shown here for the isolate *Vannella sp.* AP1411, some of the known Vannella isolates are able to form a cyst, in addition to their trophozoite stage. Moreover, Vannellidae have the particularity to harbor a star-shape pelagic form observed here for *Vannella sp.* AP1411, which may allow them to float in the water column and navigate thanks to water currents ^25^. It is likely that this planktonic form can be easily filtered by oysters while feeding, explaining why *Vannella sp.* were also found associated to oyster tissues in our survey.

We showed here that the oyster pathogen *V. tasmaniensis* LGP32 resists to predation by *Vannella sp*. AP1411. Remarkably, LGP32 is known to resist to hemocyte phagocytosis, a behavior that was shown to be central for its virulence ^17,19^. The slow growth of amoebae on LGP32 was due, to some extent, to an inefficient intracellular killing of LGP32. Indeed, after phagocytosis most intracellular LGP32 were isolated in individual phagosomes inside the amoebae without any morphological alteration whereas the non-virulent control *V. tasmaniensis* LMG20012^T^ were progressively clustered in food vacuoles and digested. This strongly suggests that LGP32 interferes with food vacuoles formation. Such an inhibition of phagosome maturation by LGP32 in amoebae is reminiscent of its capacity to inhibit phagosome acidification in oyster phagocytes ^17^.

The copper efflux p-ATPase CopA was found to play a major role in the capacity of LGP32 to resist to amoeba grazing. Particularly, CopA was shown to interfere with food vacuoles formation in amoebae, as it interferes with LGP32 intraphagosomal degradation in oyster phagocytes ^19^. As CopA is involved in the resistance of LGP32 to high concentrations of copper ^19^, our results indicate that LGP32 resistance to grazing depends on its resistance to copper concentration inside the phagosome of *Vannella sp*. AP1411. This result also indicates that intra-phagosomal copper accumulation is an ancestral highly conserved antimicrobial mechanism from free-living amoebae, social amoebae ^27^, invertebrates ^19^ to vertebrate professional phagocytes ^28^, which has applied strong selective pressures on host-microbe interactions, and that different bacterial species resisting intracellular killing have evolved copper resistance strategies to gain a fitness advantage in a diversity of predator-prey interactions.

Another important determinant of LGP32 resistance to grazing by *Vannella sp*. AP1411was the secreted metalloprotease Vsm, a major secreted protease that plays a role as a virulence factor in oysters ^20^. In our study, the Vsm deficient strain showed an increased sensitivity to grazing by the amoebae, without losing its capability to inhibit food vacuole formation intracellularly. This suggested that Vsm rather acts extracellularly rather than intracellularly during LGP32 interaction with amoebae. These observations with the LGP32 **Δ***vsm* isogenic mutant are reminiscent of the role played by other proteases secreted by vibrios in their interaction with eukaryotic cells. Particularly, the secreted protease PrtV, the InhA ortholog, participates in the resistance of *V. cholerae* to the grazing by the bacteriovorous ciliate *Tetrahymena pyriformis* ^26^. Studies on the pathogenesis of LGP32 and *Vibrio aesturianus*, another vibrio pathogenic for oysters, showed that LGP32 Vsm can induce damages to eucaryotic cells ^20^, and that its homolog Vam from *V. aesturianus*, is a virulence factor inhibiting hemocyte phagocytosis ^29^. We could then hypothesize that the secretion of the Vsm metalloprotease by LGP32 induces cytopathic or inhibitory effects on *Vannella sp*. AP1411. Accordingly, we observed striking differences of amoeba migratory speed when grazing on wild-type LGP32 or *Δvsm* isogenic mutants. Amoebae average migratory speed was doubled on top of the LGP32 ***Δ**vsm*, suggesting that Vsm disturbs grazing by affecting amoeba motility. Interestingly, recent work studying the interactions between clinical isolates of *V. cholera* and the model amoebae *Acanthamoeba castellannii* (Neff strain) reported the involvement of the HapA metalloprotease and other secreted enzymes to be involved in the resistance to predation and the fitness of *V. cholerae* during this interaction^30^.

As bacteriovorous protists exert a strong selective pressure on bacterial communities, different bacteria species are thought to have developed a diversity of survival strategies against predation through either pre-ingestional or post-ingestional adaptations ^31^. Pre-ingestional adaptions including high motility, filamentation, surface masking or toxin release are commonly found for extracellular pathogens; whereas post-ingestional adaptations including digestional resistance through vacuolar trafficking inhibition or vacuolar escaping, and toxin release are commonly found for intracellular pathogens and tend to favor bacterial growth ^31^. Here we found that, in the case of the facultative intracellular pathogen LGP32, a combination of pre-ingestional and post-ingestional adaptations could be involved. The involvement of the secreted Vsm represents a pre-ingestional adaptation, whereas the involvement of the CopA p-ATPase represents a post-ingestional adaptation, Thus, the predator-prey interactions between bacterial communities and bacteriovorus protists could represent an important driving force sustaining co-incidental selection of a diversity of virulence factors allowing some bacteria to circumvent predation by diverse of phagocytic cells from amoeba to immune cells in animals as previously hypothesized^32^. Hence such co-evolutionary dynamics may participate to the emergence of opportunistic pathogens that behave as generalists being able to colonize a diversity of hosts, like many vibrios do.

## Acknowledgments

We are grateful to Thibaut Groult, Audrey Caro, and Marc Leroy for precious help in sample collection and preparation, to Eric Abadie for field trip coordination and to Michel Cantou from the University of Montpellier for scubadiving allowing sediment collection. This work, through the use of the GENSEQ platform (http://www.labex-cemeb.org/fr/genomique-environnementale-2) from the labEx CeMEB. The authors also thank the Montpellier RIO imaging platform (https://www.mri.cnrs.fr). The present study was supported by the EU funded project VIVALDI (H2020 program, n°678589) and by the Ec2co funded Intervibrio and VibrAm projects, Ifremer, Université de Montpellier and Université de Perpignan via Domitia.

## Supplementary Figures

**Fig. S1. Phylogeny of the diversity of amoebae OTUs found by barcoding of the v7 region of the 18s rDNA gene.**

Phylogenetic 18S rRNA-based trees of the Amoebozoa constructed with ARB software (http://www.arb-home.de ^24^) loaded with the silva database (http://www.arb-silva.de). A base frequency filter was applied using the parsimony quick add marked tool implemented in ARB. All 18S rDNA gene sequences assigned to amoeba belonged to the Vannellidae order (written in red).

**Fig. S2. Quantification of amoeba phagocytosis by cytometry.**

**(A)** Gate determination corresponding to amoeba on FSC/SSC data and **(B)** Determination of fluorescence background on the PMT used to measure GFP fluorescence. **(C)** Determination of the gates of amoeba and GFP-vibrio populations in grazing experiments. **(D)** Measurement of the percentage of amoebae associated with GFP fluorescence in co-culture with GFP-vibrios.

**Fig. S3. Vsm or CopA do not affect the phagocytosis rate of LGP32 by the amoeba *Vannella* sp. AP1411.**

Grazing amoebae on GFP-vibrio LGP32wt, LGP32 ∆*copA* or of LGP32 ∆*vsm* were monitoring by flow cytometry to estimate phagocytosis index per amoeba, which correspond to the average number of phagocyted bacteria per cell. The results shown are the average of three independent experiments ± SEM. Data were analyzed by two-way ANOVA with Holm-Sidak’s multiple comparisons test.

**Description of Additional Supplementary Files**

File name: Supplementary Movie 1

Description: Time-lapse microscopy movie of *Vannella sp.* AP1411 and LGP32wt lawn in co-culture.

File name: Supplementary Movie 2

Description: Time-lapse microscopy of *Vannella sp*. AP1411 and LGP32 ∆*copA* lawn in co-culture.

File name: Supplementary Movie 3

Description: Time-lapse microscopy of *Vannella sp*. AP1411 and LGP32 ∆*vsm* lawn in co-culture.

## References

1. Lemire, A. et al. Populations, not clones, are the unit of vibrio pathogenesis in naturally infected oysters. ISME J. 1–9 (2014). doi:10.1038/ismej.2014.233

2. Bruto, M. et al. Vibrio crassostreae, a benign oyster colonizer turned into a pathogen after plasmid acquisition. ISME J. 11, 1043–1052 (2017).

3. Schmitt, P. et al. The Antimicrobial Defense of the Pacific Oyster, Crassostrea gigas. How Diversity may Compensate for Scarcity in the Regulation of Resident/Pathogenic Microflora. Front. Microbiol. 3, 160 (2012).

4. de Lorgeril, J. et al. Immune-suppression by OsHV-1 viral infection causes fatal bacteremia in Pacific oysters. doi::10.1038/s41467-018-06659-3

5. Le Roux, F. et al. The emergence of Vibrio pathogens in Europe: ecology, evolution, and pathogenesis (Paris, 11–12th March 2015). Front. Microbiol. 6, 830 (2015).

6. Constantin de Magny, G. et al. Environmental signatures associated with cholera epidemics. Proc. Natl. Acad. Sci. U. S. A. 105, 17676–81 (2008).

7. Takemura, A. F., Chien, D. M. & Polz, M. F. Associations and dynamics of Vibrionaceae in the environment, from the genus to the population level. Front. Microbiol. 5, 38 (2014).

8. Dawkins, R. & Krebs, J. R. Arms races between and within species. Proc. R. Soc. London. Ser. B, Biol. Sci. 205, 489–511 (1979).

9. Boulais, J. et al. Molecular characterization of the evolution of phagosomes. Mol. Syst. Biol. 6, 423 (2010).

10. Erken, M., Lutz, C. & McDougald, D. The Rise of Pathogens: Predation as a Factor Driving the Evolution of Human Pathogens in the Environment. Microb. Ecol. 65, 860–868 (2013).

11. Ma, A. T., McAuley, S., Pukatzki, S. & Mekalanos, J. J. Translocation of a Vibrio cholerae type VI secretion effector requires bacterial endocytosis by host cells. Cell Host Microbe 5, 234–43 (2009).

12. Ritchie, J. M. et al. Inflammation and Disintegration of Intestinal Villi in an Experimental Model for Vibrio parahaemolyticus-Induced Diarrhea. PLoS Pathog. 8, e1002593 (2012).

13. de Souza Santos, M. & Orth, K. Intracellular Vibrio parahaemolyticus Escapes the Vacuole and Establishes a Replicative Niche in the Cytosol of Epithelial Cells. MBio 5, e01506–14 (2014).

14. Rosenberg, E. & Falkovitz, L. The Vibrio shiloi/Oculina patagonica model system of coral bleaching. Annu. Rev. Microbiol. 58, 143–59 (2004).

15. Vidal-Dupiol, J. et al. Physiological responses of the scleractinian coral Pocillopora damicornis to bacterial stress from Vibrio coralliilyticus. J. Exp. Biol. 214, 1533–1545 (2011).

16. Lopez-Joven, C. et al. Oyster Farming, Temperature, and Plankton Influence the Dynamics of Pathogenic Vibrios in the Thau Lagoon. Front. Microbiol. 9, 2530 (2018).

17. Duperthuy, M. et al. Use of OmpU porins for attachment and invasion of Crassostrea gigas immune cells by the oyster pathogen Vibrio splendidus. Proc. Natl. Acad. Sci. U. S. A. 108, 2993–8 (2011).

18. Gay, M., Renault, T., Pons, A. & Le Roux, F. Two Vibrio splendidus related strains collaborate to kill Crassostrea gigas: taxonomy and host alterations. Dis. Aquat. Organ. 62, 65–74 (2004).

19. Vanhove, A. S. A. S. et al. Copper homeostasis at the host vibrio interface: Lessons from intracellular vibrio transcriptomics. Environ. Microbiol. 18, 875–888 (2016).

20. Binesse, J. et al. Metalloprotease vsm is the major determinant of toxicity for extracellular products of Vibrio splendidus. Appl. Environ. Microbiol. 74, 7108–17 (2008).

21. Vanhove, A. S. A. S. et al. Outer membrane vesicles are vehicles for the delivery of Vibrio tasmaniensis virulence factors to oyster immune cells. Environ. Microbiol. 32, (2014).

22. Wheeler, D. L. et al. Database resources of the National Center for Biotechnology Information. Nucleic Acids Res. 36, D13–D21 (2007).

23. Schloss, P. D. A High-Throughput DNA Sequence Aligner for Microbial Ecology Studies. PLoS One 4, e8230 (2009).

24. Ludwig, W. et al. ARB: a software environment for sequence data. Nucleic Acids Res. 32, 1363–71 (2004).

25. Smirnov, A. V., Nassonova, E. S., Chao, E. & Cavalier-Smith, T. Phylogeny, Evolution, and Taxonomy of Vannellid Amoebae. Protist 158, 295–324 (2007).

26. Vaitkevicius, K. et al. A Vibrio cholerae protease needed for killing of Caenorhabditis elegans has a role in protection from natural predator grazing. Proc. Natl. Acad. Sci. 103, 9280–9285 (2006).

27. Hao, X. et al. A role for copper in protozoan grazing - two billion years selecting for bacterial copper resistance. Mol. Microbiol. 00, 1–14 (2016).

28. Soldati, T. & Neyrolles, O. Mycobacteria and the intraphagosomal environment: take it with a pinch of salt(s)! Traffic 13, 1042–52 (2012).

29. Labreuche, Y. et al. Vibrio aestuarianus zinc metalloprotease causes lethality in the Pacific oyster Crassostrea gigas and impairs the host cellular immune defenses. Fish Shellfish Immunol. 29, 753–758 (2010).

30. Van der Henst, C. et al. Molecular insights into Vibrio cholerae’s intra-amoebal host-pathogen interactions. Nat. Commun. 9, 3460 (2018).

31. Matz, C. & Kjelleberg, S. Off the hook – how bacteria survive protozoan grazing. Trends Microbiol. 13, 302–307 (2005).

32. Adiba, S., Nizak, C., van Baalen, M., Denamur, E. & Depaulis, F. From grazing resistance to pathogenesis: The coincidental evolution of virulence factors. PLoS One 5, 1–10 (2010).

